# Conflict neurons in cingulate cortex of macaques

**DOI:** 10.1101/2024.10.03.616355

**Authors:** Benjamin W. Corrigan, Steven P. Errington, Amirsaman Sajad, Jeffrey D. Schall

**Affiliations:** Department of Biology, Center for Integrative & Applied Neuroscience, Center for Vision Research, Connected Minds, York University, Toronto, Canada; Biosciences Institute, Newcastle University, Newcastle Upon Tyne, United Kingdom; Department of Psychology, Vanderbilt Vision Research Center, Vanderbilt University, Nashville, Tennessee, USA

## Abstract

Conflict—the magnitude of co-activation of mutually incompatible response processes—was proposed to explain how cognitive control is invoked (Botvinick et al. 2001) and continues to engage debate (Becker et al. 2024). Original observations consistent with this construct emphasized the primary contribution of cingulate cortex (CC) based on human functional imaging (Botvinick et al., 1999; Carter et al., 2000) and electroencephalogram (Yeung, Botvinick, & Cohen, 2004). In countermanding tasks conflict arises through co-activation of competing GO and STOP processes (Boucher et al. 2007; Schall & Boucher 2007; Sajad et al. 2022). Single neuron activity representing conflict has been described in the supplementary motor cortex of human epilepsy patients (Fu et al., 2019; Sheth et al., 2012) and of macaque monkeys (Sajad, Errington, & Schall, 2022; Stuphorn, Taylor, & Schall, 2000) and in human cingulate cortex (Fu et al., 2019; Sheth et al., 2012) but not in monkey cingulate cortex (Ebitz & Platt, 2015; Ito, Stuphorn, Brown, & Schall, 2003; Nakamura, Roesch, & Olson, 2005). This lack of homology generated debate about the utility of macaques for investigation of cognitive control (Cole et al. 2009; Schall & Emeric 2010). With higher-resolution, less-biased samples, we re-examined the presence of a conflict signal in cingulate cortex of monkeys. Neurons modulating specifically when response conflict was maximal were found in cingulate cortex— more commonly in the dorsal than the ventral bank. However, such neurons were much more common in supplementary motor cortex. These data confirm the presence of a conflict signal in medial frontal cortex and demonstrate that it can be found in a small fraction of neurons in cingulate cortex. Further research is needed to determine if the weak response conflict signal in cingulate cortex is sufficient or negligible.

## Main

Two male macaque monkeys (Da, *M. radiata*, 9.5 kg and Jo, *M. mulatta*, 13.8 kg) performed a gaze countermanding (stop signal) task (**Fig. 1A**). Reward was earned for shifting gaze to a visual target unless a visual stop signal appeared, which was infrequent and a variable delay after the target. The longer the stop signal delay, the less frequently did monkeys cancel the saccade (**Fig. 1C**). Response times (RT) of non-canceled errors were consistently shorter than RT on trials with no stop signal (**Fig. 1B**). The pattern of performance satisfied the assumptions of the race model (Logan & Cowan, 1984), which enabled estimation of stop signal reaction time (SSRT), which measures when response preparation is interrupted (Boucher, Palmeri, Logan, & Schall, 2007; Logan, Yamaguchi, Schall, & Palmeri, 2015) (**Fig. 1D**) that was indistinguishable between monkeys (Da: 121 ± 2 ms; Jo: 115 ± 4 ms; Independent groups t-test, t (81) = 1.577, p = 0.119). The applicability of the race model for these data justified the measure of conflict derived from the brief co-activation of GO and STOP accumulators when responses were countermanded (**Fig. 1E**), which scaled with stop signal delay (**Fig. 1F**) (Sajad et al., 2022; J. D. Schall & Boucher, 2007). Performance of countermanding (stop signal) tasks (**Fig. 1A**) is explained as the outcome of a race between GO and STOP processes (Logan & Cowan, 1984), which can be instantiated through specific interactions between GO and STOP accumulators (**Fig. 1D**) (Boucher et al., 2007; Logan et al., 2015). The interacting GO and STOP units exhibit no co-activation on trials with no stop signal and on stop signal trials in which inhibition failed. However, when responses were inhibited after the stop signal, the interacting GO and STOP units were briefly co-active, yielded a conflict signal that scaled with how near to threshold was the GO unit when it was interrupted by the STOP unit (**Fig. 1E,F**).

**Fig. 1.**
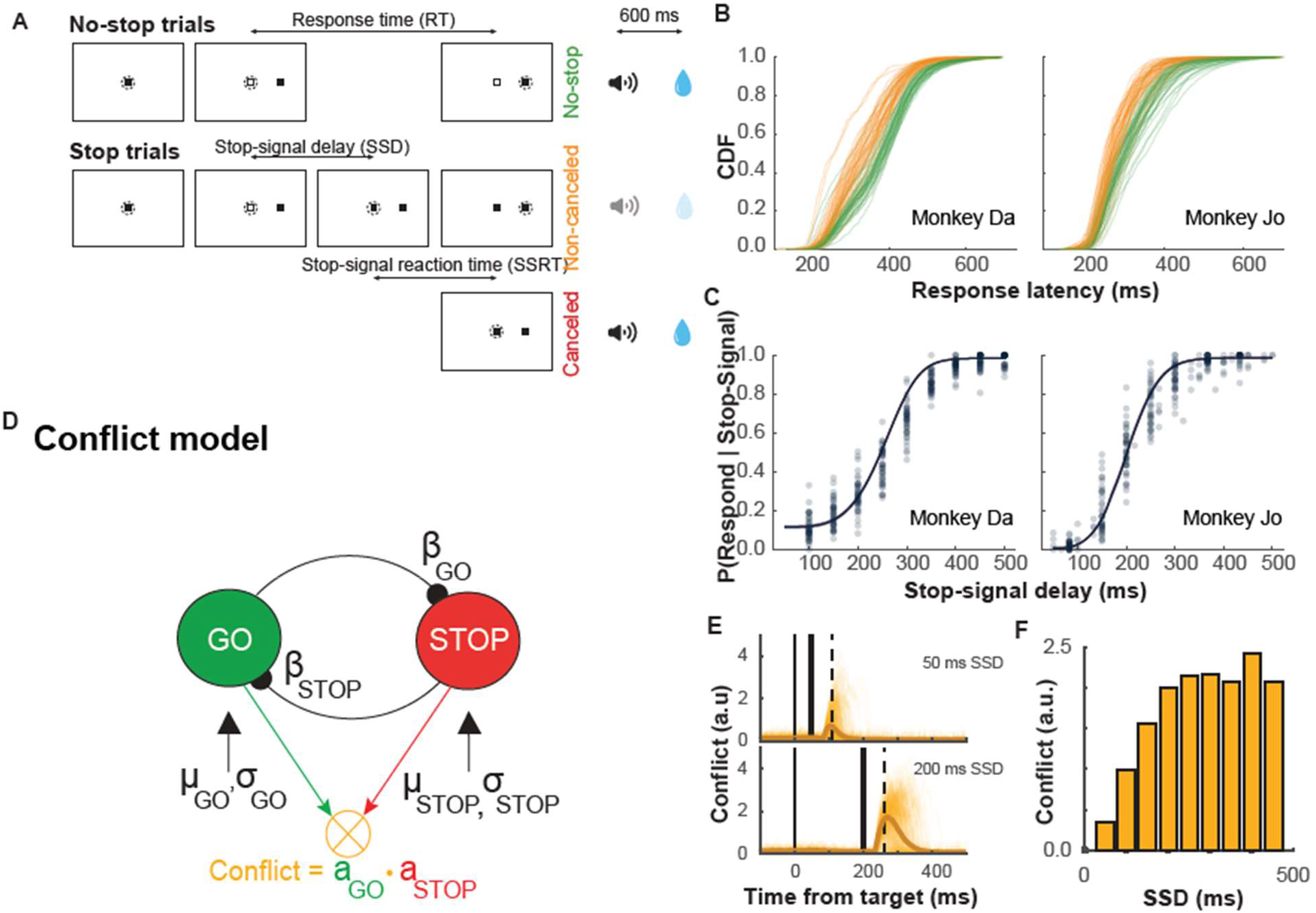
Task and Behaviour. **A)** saccade countermanding task. All trials start with central fixation, and then a target appears on either the left or the right. On no-stop trials, saccades to the target are followed 600ms later by a sound indicating correct performance and then after another 600ms, a juice reward. On stop trials, the fixation point reappears after a variable delay (SSD). Non-canceled trials where the monkey still makes the saccade are followed after 600ms by a sound indicating incorrect performance and no juice. On canceled trials, where no saccade is made, 1500 ms after the stop signal, a sound indicating correct performance plays, followed 600ms after by a juice reward. **B)** Cumulative density function of response times for non-cancelled and no-stop trials, indicating that non-cancelled responses as a population are faster than no-stop responses. **C)** Figures demonstrating that as the SSD increases, the probability to saccade on a stop trial increases. **D)** The interactive race model, where GO and STOP units are mutually inhibitory, and conflict is the product of the activation of both of them. **E)** Simulated measures of conflict for trials with SSD of 50ms and of 200 ms, where the latter is higher because the strength of the GO signal is stronger and has had longer to build towards threshold without yet crossing. **F)** Histogram of average simulated conflict values scaling with SSD (E and F adapted from Sajad et al. 2022)

During task performance, we obtained unbiased samples of neuron spiking by inserting linear electrode arrays at 34 locations 25-32 mm anterior to the interaural plane, 3-5 mm lateral to the midline, and to depths that spanned all layers of three medial frontal areas—the supplementary eye field (SEF) on the dorsomedial convexity plus the dorsal ventral banks bank of the middle cingulate sulcus (**Fig. 2A**). The dorsal and ventral banks are distinguished by connectivity and intrinsic structure (**SFig 1**) (Paxinos, Huang, & Toga, 2000; Rapan et al., 2021; Vogt, Vogt, Farber, & Bush, 2005). Being folded around the cingulate sulcus, the laminar sequence of dCC and vCC are mirror images. We located the cingulate sulcus and verified that samples spanned the cortical layers based on systematic variation of the power spectra of local field potentials measured across cortical layers (Ninomiya et al. 2015; Mendoza-Halliday et al., 2024). (**Fig. 2C**). We identified the cingulate sulcus in 94 of 104 penetrations (90%). This yielded a bimodal distribution of neuron depths with pronounced peaks approximately ±1000 um relative to the identified sulcus channel (**Fig. 2F**). In all, 2386 single neurons were sampled (813 in SEF, 956 dorsal and 617 in ventral cingulate, with 33 unassigned) (**Table 1**).

**Table 1.**
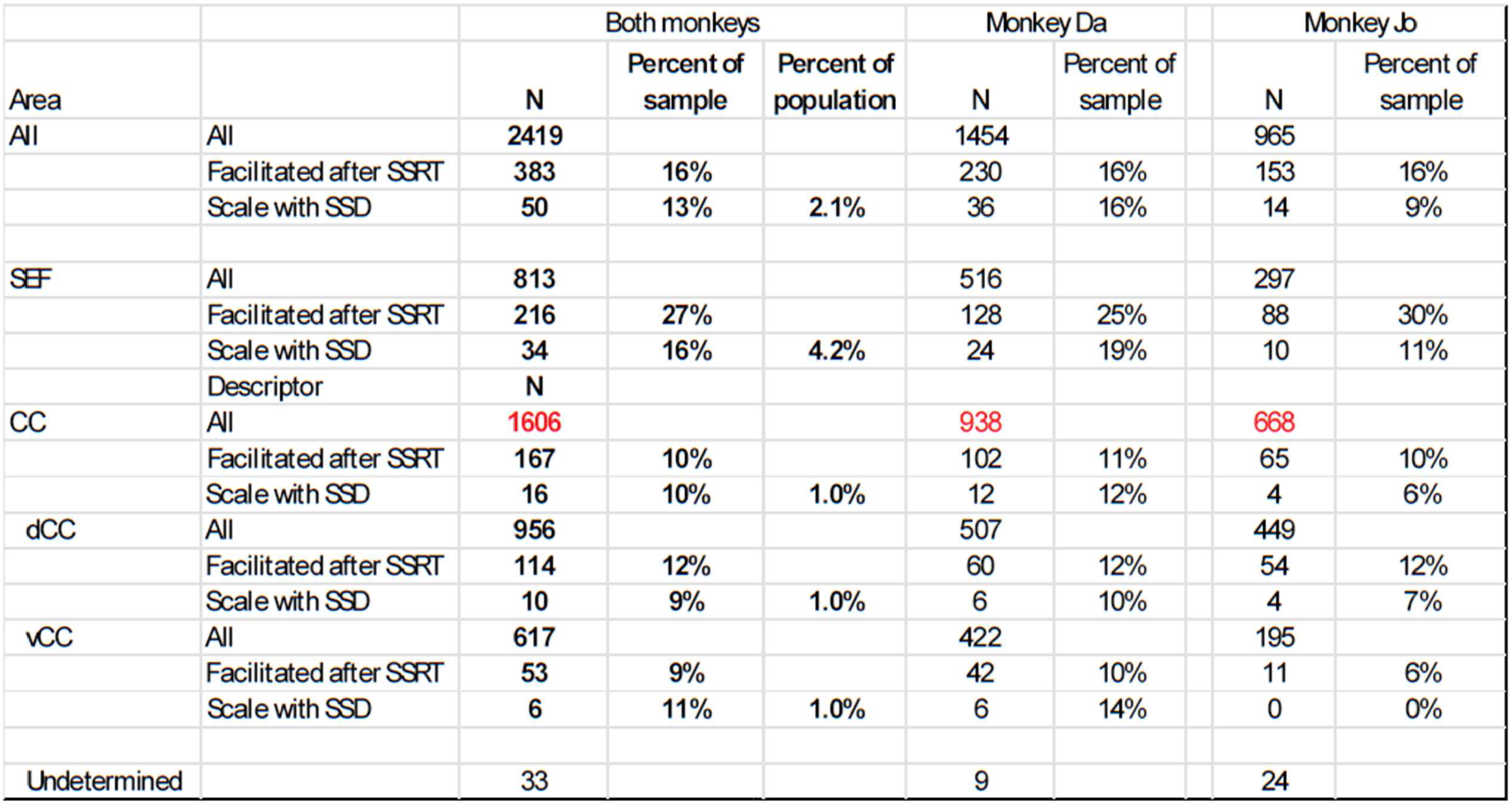
Conflict neurons in sampled population with divisions by monkey.

**Fig. 2.**
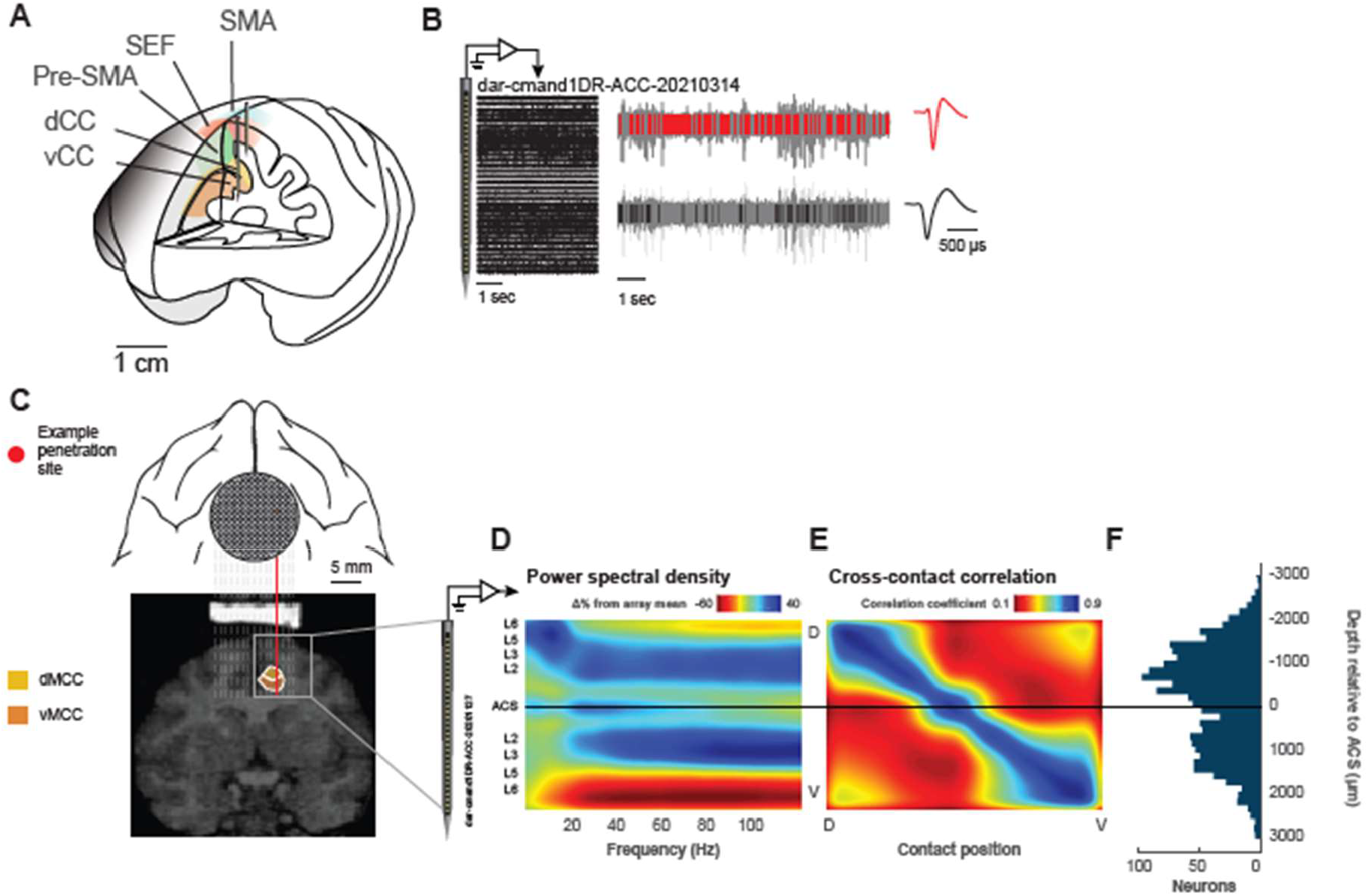
Recording locations and separation of cingulate banks. **A)** diagram of macaque brain with recording areas highlighted. The Dorsal and ventral banks of the cingulate sulcus (dCC and vCC respectively) and the supplementary motor cortex, including supplementary motor area (SMA), pre-supplementary motor area (PreSMA) and supplementary eye fields (SEF). **B)** Example of recordings across all channels and two isolated neurons from the same channel with their respective waveforms. **C)** Diagram of one chamber location with an example penetration aligned to a sagittal MRI with dCC and vCC outlined. **D)** Power spectral density across the electrodes, with higher power in the lower frequencies (<20 Hz) in the lower layers, and higher power in the gamma range (40-120 Hz) in the upper layers. The mirroring around the identified anterior cingulate sulcus (ACS). **E)** Cross contact correlations where the dorsal channels have high correlations with each other, and the same for ventral, but the correlations between ends are low. **F)** Histogram of neuron counts by depth in relation to the identified ACS channel.

To signal the presence of conflict in this task, neurons must exhibit higher discharge rates when response preparation is interrupted at SSRT (**Fig. 3**). Spike times were aligned to the SSRT estimated for each testing session for each stop signal delay (SSD) for cancelled trials and latency-matched no stop trials (RT > SSD + SSRT). Regressions of spike rate across cancelled and no stop trials were calculated in 100 ms intervals sliding in 10 ms steps. The criteria for inclusion was significantly greater (α = 0.01) activity in cancelled relative to no stop trials for at least 5 steps (**Fig. 3A,C**). We found 383 neurons that discharged when response preparation was interrupted (216 in SEF (27% of sample) and 167 (10%) in cingulate cortex) (**Fig 3, Table 1**). The difference in fractions of 17% (95% Confidence Interval = 13-19%) between SEF and cingulate cortex was significantly greater than 0%. Among cingulate cortex neurons, 114 (12%) were in the dorsal bank with 53 (9%) in the ventral. This difference of 3.3% (95% CI = 0.3-6.4%) was significantly greater than 0%. The fractions did not differ between monkeys.

**Fig. 3.**
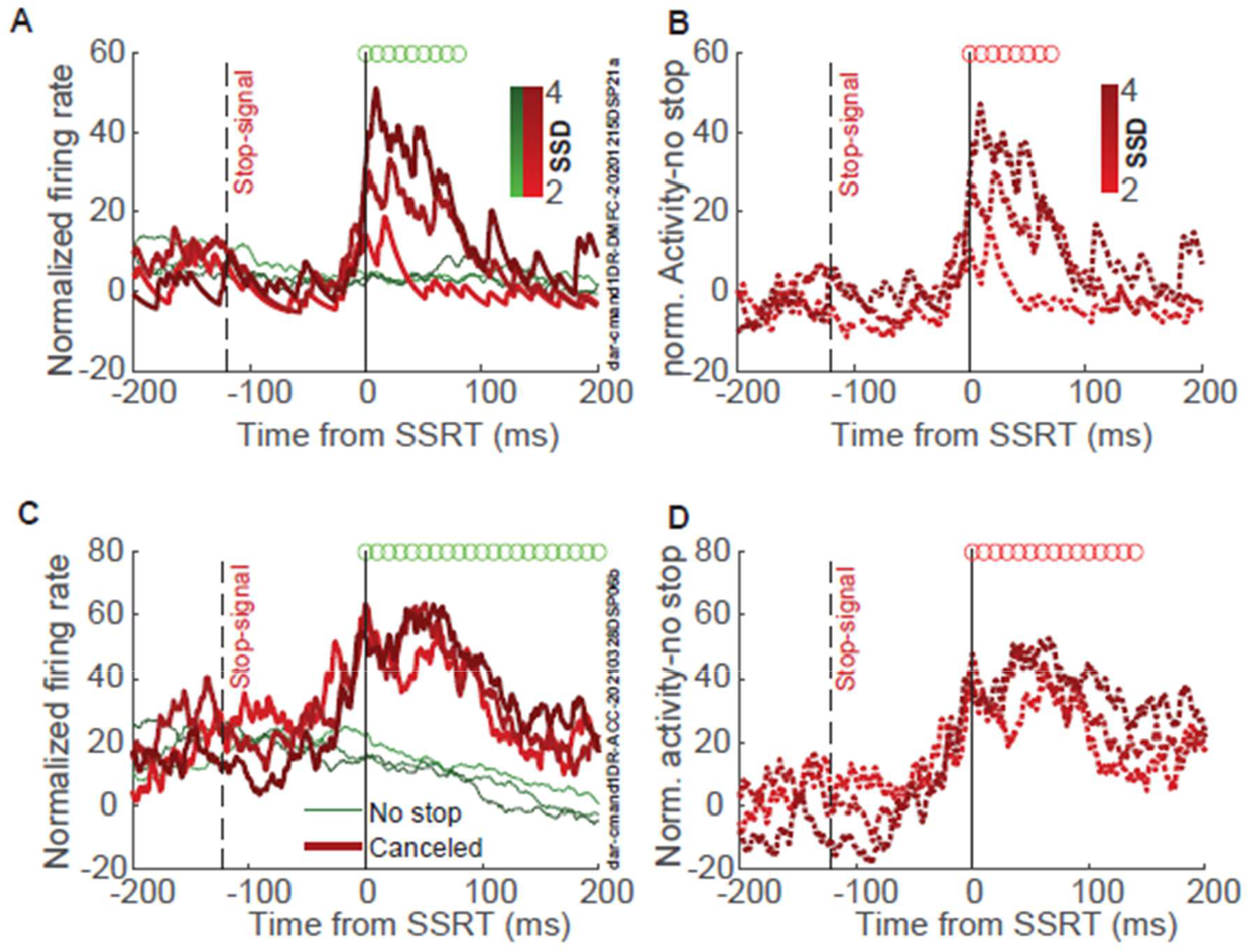
Example units signaling conflict from cingulate cortex (top) and SEF (bottom) **A**) average normalized activity of an SEF unit for cancelled (red) and latency matched no-stop trials for three different SSDs (darker = longer SSD), showing increased activity on cancelled trials, and some target related modulation pre stop signal. Green circles mark centres of time bins in which activity on cancelled trials was significantly greater than no stop. **B**) Difference in normalized activity between cancelled and no stop trials for three SSDs. Pronounced modulation happened after the stop signal and SSRT. Red circles mark centres of intervals during which cancelled activity significantly increases with SSD. **C**,**D**) same as A, B, but for CC example neuron.

To signal the magnitude of conflict, elevated discharge rate must increase with stop signal delay, paralleling the progression of the GO process to completion. Using the same regression approach to quantify whether the modulation on cancelled trials scaled with stop signal delay (**Fig. 3B,D**), we found 50 neurons that discharged when response preparation was interrupted with magnitude that increased significantly with stop signal delay (34 (16% of modulated neurons and 4% of total) in SEF and 16 (10% of modulated neurons and 1% of total) in cingulate cortex) (**Fig. 4**). The 6% (95% CI = 0.4-12.8) difference was significantly greater than 0%. Among cingulate cortex neurons, 10 (9% of modulated 1% of all) were in the dorsal bank with 6 (11% of modulated 1% of all) in the ventral. This difference of 2.5% (95% CI = -7.4 – 12.5%) was not significantly different from 0%. Sample fractions did not differ between monkeys except no scaled conflict neurons were found in vCC in monkey Jo.

**Fig. 4.**
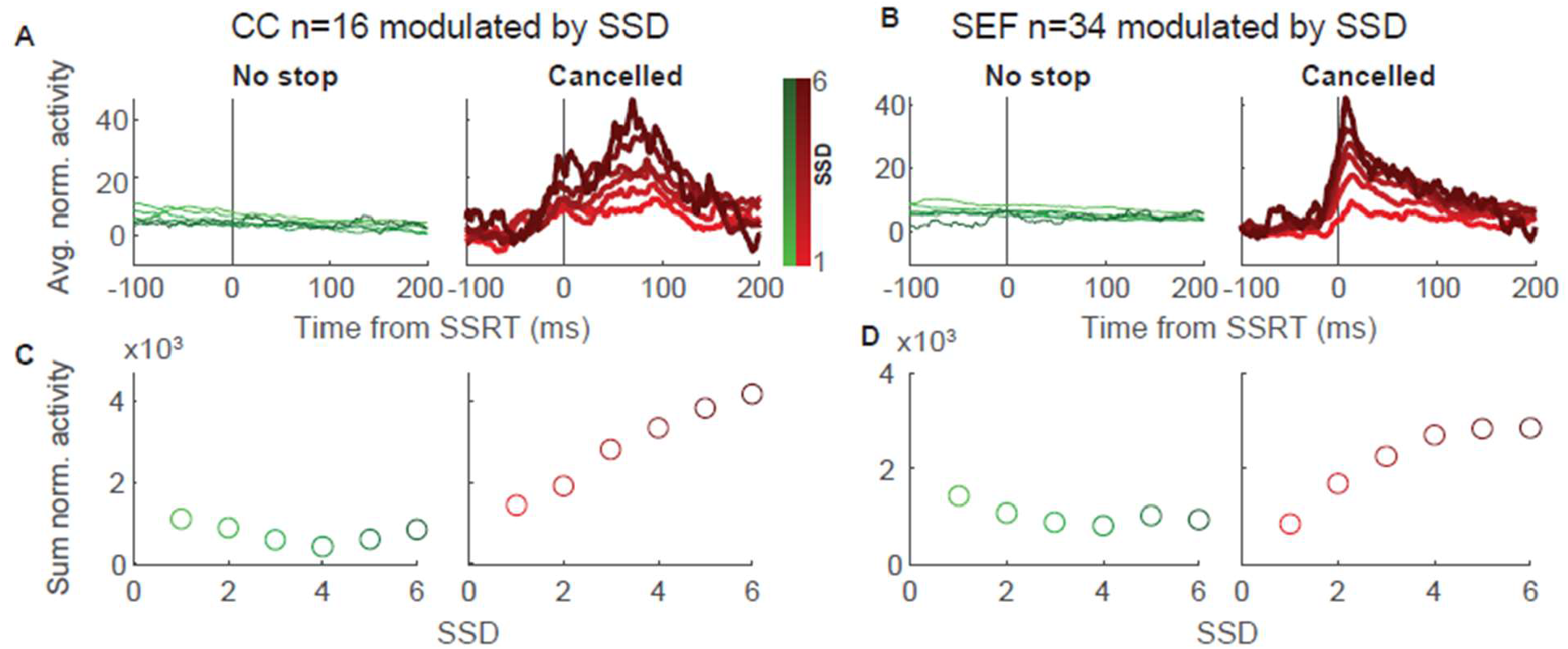
Population activity for no stop and cancelled trials for different SSDs. **A)** Activity for significant conflict scaling units in CC for no stop (left) and cancelled (right). **B**) Same as A but for SEF population. **C**) The cumulative activity from SSRT to 200ms post SSRT for no-stop (left) and cancelled (right), which has a monotonic increase. **D**) same as C, but for SEF population.

We also looked for the other combinations of facilitation or suppressions at SSRT and increasing or decreasing with SSD. While we did find similar counts of the different combinations, we wanted to look at the consistency of the effect. We regressed the cumulative sums of the average cancelled activity (200ms starting at SSRT) for the population at each SSD against SSD (**Table 2; SFig. 4**). For the cingulate conflict neurons with facilitated activity increasing with SSD the slope was 571, and there was no overlap of the 95% CI with the absolute values of any of the other populations. This suggests that the conflict neurons have a stronger and more consistent signal in CC. In the SEF, only conflict with facilitated activity increasing with SSD was significant as a population.

**Table 2.** Regressions of average cancelled activity against SSD level for different response populations. Asterisks indicate significant regressions

| Area | Canceled Activity | slope | 95% CI |
| --- | --- | --- | --- |
| CC | Facilitated Increasing * | 571 | (462 - 680) |
|  | Facilitated Decreasing | -25 | (-249 - 199) |
|  | Suppressed Increasing * | 180 | (109 - 251) |
|  | Suppressed Decreasing * | -178 | (-295 - -61) |
| SEF | Facilitated Increasing * | 398 | (185 - 611) |
|  | Facilitated Decreasing | 46 | (-385 - 477) |
|  | Suppressed Increasing | 62 | (-58 - 182) |
|  | Suppressed Decreasing | -186 | (-369 - -7) |

## Discussion

Here we described the presence of conflict neurons in SEF and in both banks of the CC in monkeys carrying out a saccade countermanding task. The finding in CC is important because there have been a few studies that have looked for action conflict facilitated neurons in the macaque mid cingulate region that did not find them (Ebitz & Platt, 2015; Ito et al., 2003; Michelet et al., 2016; Nakamura et al., 2005). Action conflict, as defined by Botvinick and colleagues (2001) requires that there be different action plans that are incompatible, and so the saccade countermanding task achieves this by first cueing a saccade, and then the stop signal indicates that there should be no saccade, creating conflict between these two motor plans. Ito and colleagues used a countermanding task, and sampled 454 neurons, mostly form the dorsal bank of the sulcus. They did not find any neurons that had differing modulation between cancelled and latency matched no-stop trials (Ito et al., 2003). Considering that, our results are surprising, as we found 10% of our larger population at least had that type of modulation. One explanation, in addition to the smaller sample size, could be that they used a fairly conservative threshold of a difference of 6 times the standard deviation of the baseline period to be detected. While our methods are still quite conservative, they do not require such a magnitude of activity change, which could be why we were able to detect them.

Nakamura and colleagues used a pro- and anti-saccade task, and also combined it with a change signal task where a pro-saccade signal could be changed to an anti-saccade, or vice versa (Nakamura et al., 2005). They do not report the total number of recorded neurons, but instead only report on neurons that were modulated during the peri-cue or peri-saccade period; 172 for the pro- and anti-saccade task, and 144 for the combination task. They also reported on SEF recordings and found mixed-selectivity for conflict and saccade direction there, but did not find more conflict neurons in their small cingulate population than expected by chance. They used ANOVAs, and so should have had the same capacity to detect activity as in our approach, but it could be that the levels of action conflict were too low to detect. While they reported behavioural effects on errors and reaction times, the tasks they used either had each action plan building up almost simultaneously in the first task, or the instruction changed after only 100ms in the second task. For the first task, there is no time for one action plan to build up as in the conflict modeled with the interactive race model (**Fig. 1D**) (Boucher et al., 2007; Sajad et al., 2022), and so that could be why there was no detected conflict in CC. For the second task, 100ms may still not be enough time for conflicting action plans to build up enough activity to be detected by conflict neurons in CC. The countermanding task on the other hand does allow for the signal to build up; we only used one SSD that was below 100ms per session, and non-cancelled probability does not really start increasing until SSDs of 150 ms (**Fig.1C**). Additionally, because there was a fixed interval for the change signal in the Nakamura task, it could be that the first motor plan wasn’t strongly invested in until the second 100ms, again resulting in limited conflict between action plans. The simultaneity argument during anti-saccades could also explain why action conflict cells were not found in CC in the social interference task (Ebitz & Platt, 2015). It should be noted that they do describe task conflict neurons, cells that increase their firing rate when a distractor was either congruent or incongruent compared to the neutral position.

Michelet and colleagues used a Stroop-like task where monkeys associated a shape with a colour, and then the shape could either be filled with white, the congruent colour, or an incongruent colour. They found that response time patterns across trial types were similar to humans doing the same task and doing the actual Stroop task (Michelet et al., 2016). They did report that there were cells in CC that responded on incongruent correct trials, which they note could be monitoring conflict, but conclude that because there is also conflict on incongruent error trials, and none of the cells were also facilitated on these trials, that these cells couldn’t be conflict cells. In contrast, an example of error conflict activation can be seen in SFig. 2, where activation is not as high as in cancelled trials but has a similar pattern. One explanation for their lack of finding this activity could again be that the action plans should have been simultaneously generated, and so perhaps on the error trials there just was not a second correct action plan to generate conflict. If incorrect incongruent trials were faster than correct, and therefore more similar to the congruent response times, this might be a reasonable proposal, however they did not report the incongruent error reaction times. Additionally, this task didn’t have multiple levels of conflict, so they were unable to examine scaling of response. Finally, they also only recorded from 403 neurons, and so just might have not recorded from action conflict neurons.

Action conflict units have been described in monkeys in SEF (Sajad et al., 2022; Stuphorn et al., 2000) as well as in dopamine cells in the ventral midbrain (ventral tegmental area and substantia nigra pars compacta) (Ogasawara, Nejime, Takada, & Matsumoto, 2018). Based on the subpopulation of the targeted dopamine neurons, Ogasawara and colleagues found that 37% of cells detected the presence of conflict, and as a population, they scaled activity with SSD (Ogasawara et al., 2018). In SEF, smaller populations of conflict presence cells were described: 13% in both Stuphorn et al. (2000) and Sajad et al. (2022). In both studies these cells scaled activity with SSD at the population level. We found a higher proportion of conflict presence cells (27%), but only 16% of them individually significantly scaled with SSD (4% of the total population). While the first population does appear larger, we did not use microstimulation to verify the location of SEF and could have recorded from the wider supplementary motor cortex than just SEF, where incidence might be higher. The second population looks smaller, but these neurons each individually scaled with SSD, analyses not reported in the other studies.

Moving to compare our results with human studies, the species where conflict activity in the cingulate was detected first on fMRI (Botvinick et al., 2001; Carter et al., 1998), we can examine results from epilepsy patients completing tasks that generate conflict. Sheth and colleagues first described conflict cells using the multi-source interference task (MSIT) (Sheth et al., 2012). They not only found units that increased activity on trials with conflict, but by coding the magnitude of conflict as whether there were one or two sources of interference, they also found units that had activity that scaled with conflict, largely in the period between cue and response. In contrast, our results for both CC and SEF are mostly after the SSRT, when activity cannot contribute to the current trial response. However, they only show an example neuron, and then analyse population responses based on when the strongest response was, but it should be noted that this population of 24 units was ∼40% of the total population. Whether the whole population are conflict responsive or just on average is not clear. In contrast, Fu and colleagues also reported the presence of conflict units during a Stroop task (Fu et al., 2019). However, they used regression on each neuron and found only ∼6% of cingulate neurons increased activity during trials with conflict, which is much closer to the 10% that increased activity on cancelled trials in this report. They did not report on conflict scaling, so we cannot compare, but it does look like the more conservative approach of analysing for the effect of conflict specifically does yield small populations of conflict detecting cells in both species in cingulate cortex.

Another difference between previous studies and the current one is that the original investigation of cingulate cortex in monkeys performing the countermanding task detected neuron spiking with single sharp electrodes, which yielded relatively small and likely biased samples. The advent of linear electrode arrays that can be positioned to sample neuron spiking across all cortical layers offers an opportunity to obtain a larger and less biased sample. It was using these that we sampled from both banks of the cingulate and SEF and found conflict sensitive and conflict scaling units in all three regions. The size of the conflict scaling population was larger in SEF, where these cells had been described before, but these were all small populations. Future experiments that make use of even higher density electrode arrays will give higher yields and hopefully shed even more light on the properties of these neurons.

An important clarification about the small populations is that while we used alphas of .01, a population of only 1% is not simply the false discovery rate because we ran two consecutive analyses that are not dependent on each other. Just because activity increases with SSD on cancelled trials does not inherently mean that the average cancelled activity must also be greater than activity on no-stop trials. Therefore, the proper percentage to compare to the alpha is the percentage of the units that responded to the presence of conflict (10% in CC and 16% in SEF). We also ran the same analyses looking for the other combinations of facilitation or suppression on cancelled trials and increasing or decreasing firing rates with SSD (**SFig. 4**). In both cingulate cortex and SEF, the population of facilitated conflict neurons (facilitated and increasing) had the strongest relationship between SSD and firing rate compared to other responses to conflict presence (suppressed) and SSD (increasing or decreasing) (**Table 2**).

These results, combined with the findings in both humans and monkeys, suggests that there is a population of neurons that respond to the presence of conflict, but not all of them respond to the magnitude. Depending on the composition of the sample, it might be the case that the population average is significantly modulated by magnitude, but not guaranteed. Now, These analyses are all focused on which computations about a specific component of the task neurons are computing, and is anti-thetical to current trends of analysing population codes and mixed selectivity (Diester et al., 2011; Eichenbaum, 2018; Fusi, Miller, & Rigotti, 2016). Our approach is supported by previous work in SEF, which showed that neurons that encode conflict are not the same neurons that encode errors (Fu et al., 2019; Sajad, Godlove, & Schall, 2019; Stuphorn et al., 2000). These types of analyses are what support hypothesizing circuits that could compute the different calculations that cognitive control requires (Fu, Sajad, Errington, Schall, & Rutishauser, 2023; Sajad et al., 2019). However, the proposed circuits from previous papers do require that information converge at some point, which suggests there should be mixed selectivity cells, and indeed, Nakamura and colleagues (Nakamura et al., 2005) did report mixed-selectivity for conflict and saccade direction in SEF. Given the small populations of conflict cells reported here, and the importance of conflict magnitude in the models of Botvinick et al. (2001) there could well be cells that are mixed selective in CC and SEF that our methods did not detect. However, analysing for potential mixed selectivity would have extended beyond the scope of the current study, which is to simply report the detection of conflict units around the cingulate sulcus of macaques.

This brings us to another pertinent question: whether the dorsal bank is an extension of supplementary motor area (Rapan et al., 2021; Vogt et al., 2005) or a transition zone between the adjacent areas (Paxinos et al., 2000)? Our goal is to simply report differences between the two banks, and so identifying the sulcal separation between the two is important. In terms of those findings, the critical factor in this report is that there are conflict units in dCC. The sampling in vCC was much lower, and lowest in the animal that we did not find any scaling conflict units in, so their presence there is debateable, but cells that were facilitated on conflict trials were observed here.

The presence of these neurons in both humans and macaques suggests that, in contrast to Cole et al. (2009) macaques are a good model species for studying cognitive control (Jeffrey D. Schall & Emeric, 2010). This homology of the medial frontal cortex between monkeys and humans is important because of how many diseases and disorders can disrupt cognitive control, including schizophrenia (Thakkar, Schall, Heckers, & Park, 2015), ADHD (Armstrong & Munoz, 2003; Schachar, 2023) and OCD (Penadés et al., 2007). Having a model with which we can test the effects of cognitive control treatments such as pharmaceuticals or stimulation will drastically facilitate their development. It is also possible that cortical layer information could be leveraged to target specific parts of the cortical microcircuit instead of inundating the whole area with stimulation. While perhaps the cells that we report are too few to confidently ascribe to belonging to a specific layer (**SFig. 3**), knowing whether human and macaque circuits contain the same component neurons, which are likely carrying out the same computations, means that other parts of the circuits might be targeted, such as those engaged in error processing (Fu et al., 2023, 2019; Herrera, Sajad, Errington, Schall, & Riera, 2023; Sajad et al., 2019).

The existence of action conflict detection units in cingulate cortex was postulated over 20 years ago (Botvinick et al., 2001), and this theory of CC function, that it monitors performance and the presence and magnitude of conflict in order to guide cognitive control still engages the scientific community (Becker et al., 2024; Botvinick, Cohen, & Carter, 2004; Fu et al., 2022; Sheth et al., 2012). This study reinforces this theory by describing the hypothesized neuronal activity in a second species. Future work to better characterize the circuits within the areas and the network between areas will give a better understanding of the contributions of these populations of neurons. We found more neurons in SEF and the surrounding area, and some studies have actually placed the conflict activity more in pre-supplementary motor area rather than CC in humans (Hester, Fassbender, & Garavan, 2004). Simultaneous recordings in different regions could give a better understanding of how information flows between these areas, and how they could work together to facilitate cognitive control.

## Methods

### Experimental subjects

Data was collected from one male bonnet macaque (*Macaca radiata*, Da, 9.5kg) and one male rhesus macaque (*Macaca mulatta*, Jo,13.8kg). They preformed a saccade countermanding task (Hanes & Schall, 1995). All procedures were approved by the Vanderbilt Institutional Animal Care and Use Committee in accordance with the United States Department of Agriculture and Public Health Service Policy on Humane Care and Use of Laboratory Animals.

### Task

Data was recorded from monkeys performing a saccade countermanding task (Hanes and Schall, 1995; Godlove et al., 2014). The saccade stop-signal task utilised in this study has been widely used previously (Hanes & Schall 1995). Recently, a set of guidelines has been proposed for designing and analysing the stop-signal task to allow for valid comparisons to be made across studies (Verbruggen et al., 2019). Our study followed all of the recommendations but two. These adjustments were necessary to obtain sufficient neural data and to address issues arising because monkeys gain so much more experience with the task parameters relative to human participants. First, 30-40% of trials in our study were stop trials, compared to the recommended 25%. This higher value was used to achieve the necessary power to analyse neural data at the individual stop-signal level, but it did not introduce excessive slowing of responses (Emeric et al., 2007). Secondly, although we employed a staircase procedure, this stepped between one to three stop-signal delays. We do this to prevent monkeys from anticipating the staircase (Nelson et al., 2010).

Saccade countermanding tasks for macaques have been previously described (Godlove et al., 2011; Hanes & Schall, 1995; Sajad et al., 2019). Briefly, the monkey initiated a trial by fixating on a central point (Fig. 1). After a non-aging foreperiod, a target appeared peripherally either to the right or left of fixation, and simultaneously, the centre of the fixation point extinguished, leaving only an outline. On the majority of trials, the monkey was rewarded for saccading to the target (no-stop trials). However, a minority of trials were stop-signal trials, and on these trials after a short delay, the fixation point would reappear, which was the stop-signal (stop-signal trials). The stop signal indicated that the monkey should cancel the planned fixation and maintain fixation at the centre. The delay before the stop signal is called the stop signal delay (SSD). If monkeys maintained fixation on stop trials, they received a reward and did not receive a reward if they broke fixation, even if they did so before the appearance of the stop signal. The range of SSDs was adjusted to ensure that each monkey failed to countermand the saccade on ∼50% of stop-signal trials by using a staircase procedure to lengthen the SSD after failures to cancel saccades and shorten it after successful cancellations. Following each completed trial was a 600 ms delay and then an auditory tone was played, one for successful trials, and another for incorrect trials. After another 600 ms delay, correct trials were followed by delivery of a fluid reward.

### Surgical procedures

Surgical details have been described previously (Godlove et al., 2011). To guide chamber positioning, Magnetic Resonance Images (MRIs) and have been described previously (Thirunavukkarasu et al., 2023). The chamber for monkey Da was implanted over both hemispheres, normal to the cortex, centred on the midline, 30 mm anterior to the interaural line. The chamber for monkey Jo was implanted over the left hemisphere, normal to the cortex (28° relative to stereotaxic horizontal, 19° relative to stereotaxic vertical), -1.2 mm lateral to the midline, and 33.3 mm anterior to the interaural line.

### Electrode Placement

Neural spiking and local field potentials were sampled using linear electrode arrays (NeuroNexus, 32 channels with 50, 100, or 150 µm spacing; Plexon, 32 channels with 100 or 150 µm spacing). Recordings were acquired from both the dorsal and ventral banks of the cingulate sulcus (dCC and vCC respectively) as well as from SEF. From monkey Da, cingulate cortex samples were acquired in 47 penetrations located between 26 and 34 mm anterior to the interaural line and 3 to 5 mm lateral to the midline. SEF samples were obtained from 32 penetrations located 27-33 anterior to the interaural line and 2-6 mm lateral to the midline. From monkey Jo, cingulate cortex samples were acquired in 48 penetrations 28 to 33 mm anterior to the interaural line, 4 to 6 mm lateral to the midline. SEF samples were obtained in 24 penetrations 28-37.5 mm anterior to the interaural line, 3-6.5 mm lateral to the midline.

### Data collection protocol

An identical daily recording protocol across monkeys and sessions was carried out. In each session, the monkey sat in an enclosed primate chair with their head restrained 57 cm from a CRT monitor (NEC MultiSync FE922-BK). The monitor had a refresh rate of 60 Hz, and the screen subtended 37° x 30° of the visual angle. Eye position data was collected at 1 kHz using an EyeLink 1000 infrared eye-tracking system (SR Research, Kanata, Ontario, Canada). Behavioral and neural data was streamed to a single data acquisition system (TDT System 3, Tucker-Davis Technologies, Alachua, FL). Time stamps of trial events, eye position, photodiode signals, and reward solenoid signals were recorded at 2034.5052 Hz.

### Electrophysiological recordings

Intracortical spiking activity and local field potentials were recorded using 32-channel NeuroNexus acute vector arrays (Ann Arbor, MI) with spacings of 50, 100 or 150 µm and Plexon S-probes (Dallas, TX) with spacings of either 100 or 150 µm. The spacing allowed for unbiased sampling across cortical areas. Both Arrays and S-Probes were housed in custom built guide tubes made from 21-guage stainless steel hypodermic tubing (Small Parts Inc., Logansport, IN). Tubing was cut to length, deburred and polished to effectively support the electrodes as they penetrated dura and entered the cortex. For both intracortical electrodes, contacts were referenced to the guide tube in which they were housed and grounded to the headpost.

Microdrive adapters were fit to recording chambers with < 400 µm of tolerance and locked in place at a single radial orientation (Crist Instruments, Hagerstown, MD). After setting up hydraulic microdrives (FHC, Bowdoin, ME) to move the adapters, pivot points were locked in place by means of a custom mechanical clamp. Guide tubes and electrodes remained attached to the microdrives for the duration of recording. These methods ensured that we sampled neural activity from precisely the same location relative to the chamber on repeated sessions.

Electrodes were connected to the recording system through high-input impedance ZIF-clip head stages (ZD32, Tucker-David Technologies), and signals digitised through a preamplifier (PZ5, Tucker-David Technologies) and streamed to a single data acquisition system (TDT System 3, Tucker-Davis Technologies, Alachua, FL) at 24,414 Hz. Neural data was filtered online through the software Synapse (Tucker-Davis Technology). Local field potentials were bandpass filtered between 3 and 300 Hz, with notch filters at 60Hz, 120Hz, and 180Hz. Spiking data were derived from a broadband signal, bandpass filtered between 300 and 5000 Hz. Single units were extracted offline, identified using automated spike sorting via Kilosort, and manually curated thereafter.

### Cortical depth assignment

Depth of CC units was assigned based on distance to a specific channel identified as being in the cingulate sulcus. This assignment was done manually based on the power spectral density modulation that should be mirrored at the cingulate sulcus (**Fig. 2D**) and cross contact correlation that should measure similar values within a bank and different values across the bank. We also used the assigned sulcus channel as a 0 point, and calculated depth in relation to that to measure unit clustering across sessions.

### Conflict assessment

To determine whether a unit was significantly modulated by conflict, we aligned trial activity on cancelled trials to the SSRT for that session at each SSD. To compare to no stop trials, for each SSD we selected a matching number of no stop trials that had reaction times that were later than the particular SSD + SSRT. Selected trials were removed from the pool of potential matches for other SSDs to prevent double sampling. We then calculated firing rates over 100ms bins starting at 50ms before SSRT and ran a regression with betas for cancelling and normalized trial time to determine if cancelled activity was higher than no stop activity. We used a permutation test with 500 permutations to determine if the calculated beta for cancelling was greater than 99% of the permutations. We then moved the window in 10 ms steps and repeated the process until the window started at 250ms. We only considered a unit selective if it was significant for at least 5 consecutive steps.

To determine whether the neuron activity tracked the magnitude of the conflict, we again ran a regression, this time only on cancelled trials using SSD level and normalized trial start time as regressors. However, because there could be influences of the target appearance that are shifted in time based on the SSD, instead of using the raw firing rates, we subtracted the mean latency matched no-stop activity from the activity on each cancelled trial. We used the same permutation methods and only chose neurons that had coefficients for SSD that were greater than 99% of the permutations for 5 consecutive windows. These windows needed to be the same windows during which the neuron was selective for cancelling.

We repeated this analysis for other activity patterns that were the remaining combinations of suppressed or facilitated by SSRT and increasing or decreasing with SSD. We took the cumulative sum of normalized activity for each SSD level from one to six and ran a regression on the values. Each neuron only contributed to an SSD where there were at least 7 canceled trials in that session. To determine how consistent the neurons in the population were, we calculated the 95% confidence intervals and used those to determine if the regression was significant, and if there were significant differences between populations in terms of the consistency of the average activity.

## ACKNOWLEDGEMENTS

This work was supported by the National Institute of Mental Health (R01MH55806); National Eye Institute (P30EY008126); Canadian Institutes of Health Research Postdoctoral Fellowship; and Natural Sciences and Engineering Research Council of Canada (RGPIN-2022-04592).

## Data availability

Data available on request.

## Code availability

Code available on request.

## Supplementary Figures

**SFig 1.**
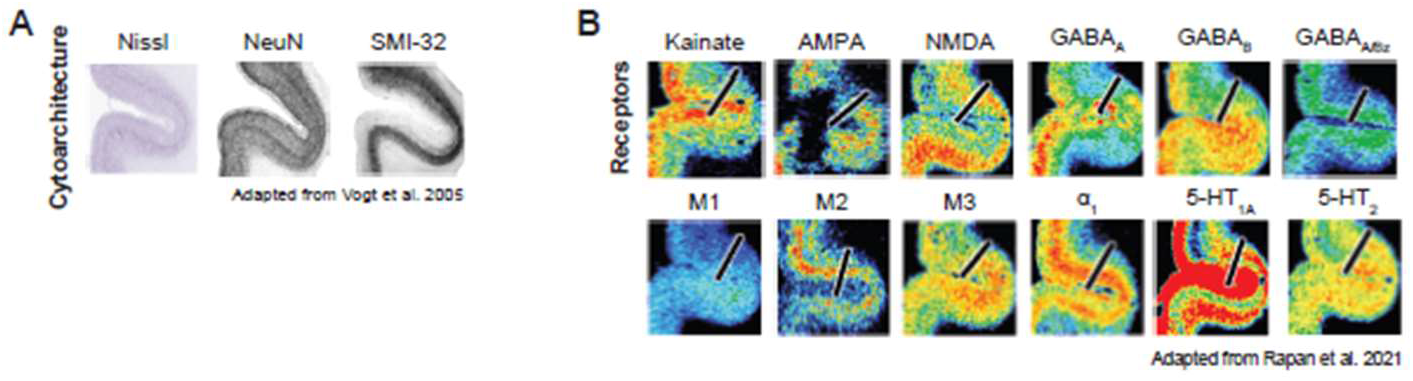
Histology depicting the separation between dorsal and ventral cingulate banks. **A)** Cytoarchitecture stains, adapted from Vogt et al., 2005. **B)** receptor stains where the border between the regions is depicted with a black line. Adapted from Rapan et al., 2021.

**SFig 2.**
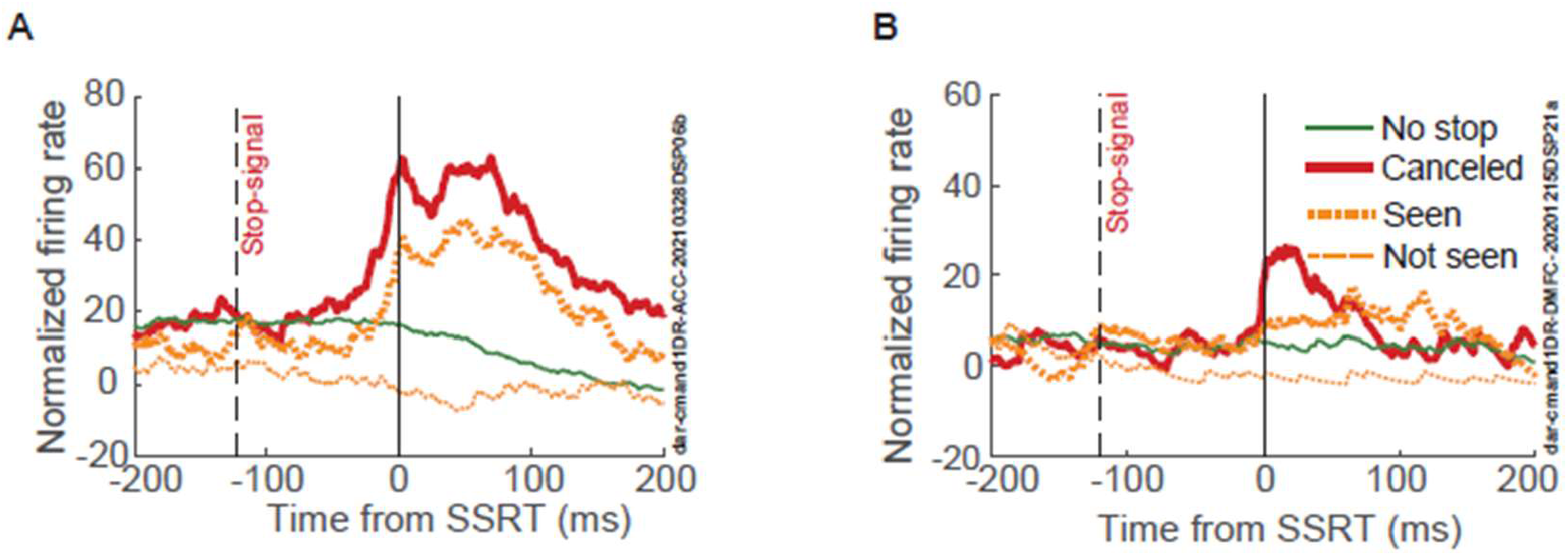
Conflict activity is higher than seen uncanceled activity. **A)** Activity aligned to SSRT for cancelled trials, latency matched no stop trials, and non-canceled trials where the stop signal was seen (Seen) and where the saccade occurred before the stop signal (Not seen).

**SFig. 3.**
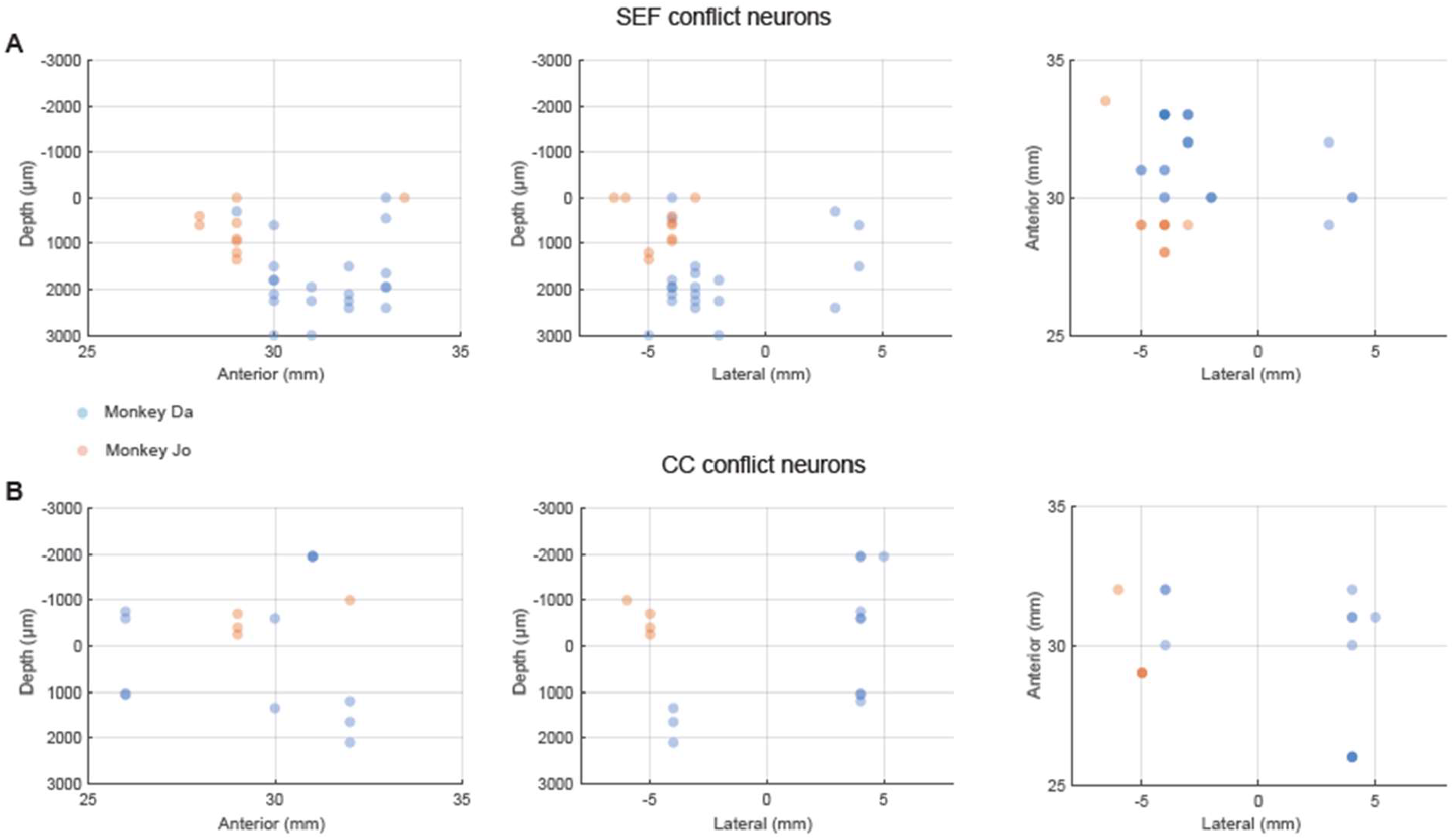
Locations of conflict neurons. **A)** Locations of CC electrode contact where neuron was recorded in relation to the identified sulcus channel (0), anterior to the interaural plane, and lateral from the midline for each monkey. **B)** Locations of SEF electrode contacts where neurons were recorded in terms of depth from the top of cortex, anterior to the interaural plane, and lateral from the midline for each monkey.

**SFig. 4.**
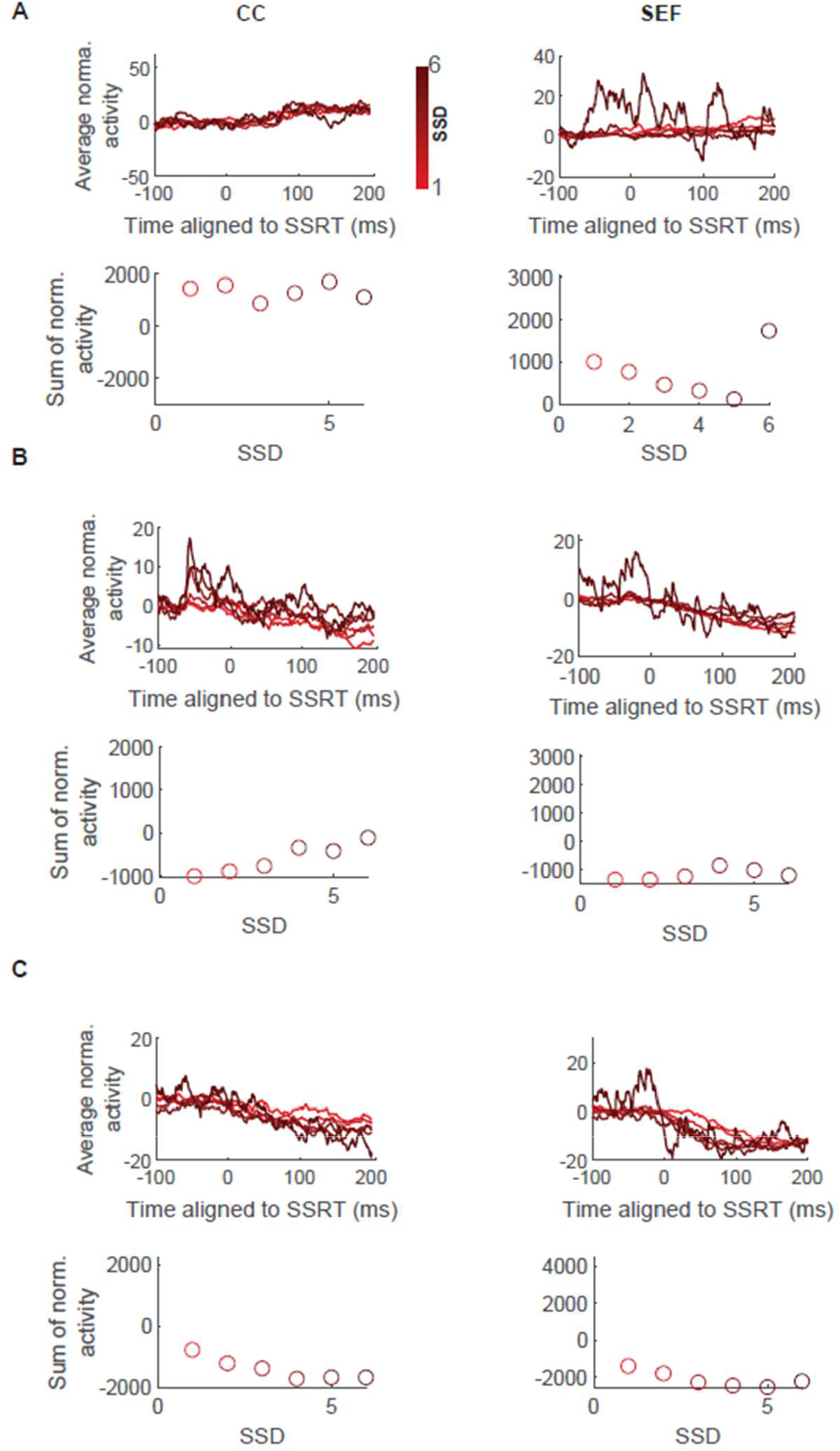
Average population responses for other combinations of SSRT modulation and SSD scaling. **A)** Average activity for neurons significantly facilitated by SSRT and decreasing with SSD, and cumulative sum of 200ms of activity starting at SSRT. **B)** Average activity for neurons significantly suppressed by SSRT and increasing with SSD, and cumulative sum of 200ms of activity starting at SSRT. **C)** Average activity for neurons significantly suppressed by SSRT and decreasing with SSD, and cumulative sum of 200ms of activity starting at SSRT.

